# The NMDAR positive allosteric modulator NYX-783 selectively blocks opioid withdrawal conditioned place aversion in mice

**DOI:** 10.1101/2025.09.25.678598

**Authors:** Joseph R. Trinko, Ethan Foscue, Deven M. Diaz, Summer L. Thompson, Robert T. Malison, Rajita Sinha, Jane R. Taylor, Ralph J. DiLeone

## Abstract

Rates of opioid use disorder (OUD) and overdose deaths have increased dramatically in recent years. Currently approved medications for OUD include the opioid agonists methadone and buprenorphine, and the opioid antagonist naltrexone. However, relapse rates are high due to an impaired ability for addicts to control their urge to consume due to strong cravings, and the extreme severity of withdrawal symptoms. Additionally, the nature of the agonists can lead to abuse of those compounds as well. Non-opioid targets, such as glutamate receptors, are potentially ideal for developing intervention strategies with the goal of reducing OUD, relapse, and overdose death. NYX-783 is a small molecule positive allosteric modulator for the glutamate receptor NMDA. It has been shown to modulate learning and memory, both of which are impaired in drug addicts, and known play a role in relapse. Using mice, we have conducted preclinical studies to evaluate the potential for NYX-783 as a therapeutic for OUD with assessment of several outcomes: 1) respiratory depression, 2) consumption during maintenance of regular consumption as well as post-abstinence reinstatement, 3) motivation for consumption, 4) somatic withdrawals, and 5) the development of aversion to withdrawal symptoms. For respiratory depression studies, mice were pretreated 1h prior with NYX-783 and respiratory rates were monitored for 15min after each escalating dose of oxycodone. No effects were seen for any dose of NYX-783. To test for effects on drug consumption, mice were trained to orally self-administer oxycodone and then treated every two days with different doses of NYX-783 or entering an abstinence phase prior to testing with NYX-783. No effect on intake at clinically-relevant doses was observed during regular maintenance or post-abstinence intake. Additionally, we used a progressive ratio to assess the motivation self-administering mice had for reward acquisition, and no effect of NYX-783 was observed on rewards earned. We next evaluated withdrawal using two separate paradigms to test for effects of NYX-783 on, 1) for somatic withdrawal symptoms, 2) aversion to the state of withdrawal. For somatic withdrawal, using higher doses of naloxone (1 mg/kg), NYX-783 did not attenuate jumping behavior. For aversion to withdrawal, three aversion pairings were completed that consisted of oxycodone treatment, followed by NYX-783 preceding a low dose of naloxone (0.1mg/kg ip) immediately before pairing in a specific context. These alternated with neutral (saline) pairings daily. We observed a significant improvement in aversion scores in female mice treated with NYX-783, and a trending significant improvement in males. This suggests a potential therapeutic use for NYX-783 in reducing the negative state of withdrawal that can drive relapse in OUD.

## Introduction

Opioid use disorder (OUD) is a major public health crisis. Overdose death rates increased by more than 500% between 1999 and 2019, and 8.9 million people aged 12 or older misused opiates in 2023^1^. While prescription opioids such as oxycodone are often used for pain management, this can lead to misuse and transition to illicit opioid abuse. In 2022 there were 79,358 opioid-related deaths in the United States^2^. Despite the scale of the crisis, there are currently only three FDA-approved medications for the treatment of OUD: methadone and buprenorphine, both of which are mu-opioid receptor agonists, and extended-release naltrexone, a mu-opioid receptor antagonist^3^. The majority of patients who exclusively undergo outpatient medical detoxification typically relapse at a high rate^4^. While longer-term medical interventions with buprenorphine-naltrexone combinations have demonstrated an improvement in patient outcomes over monotherapy, retention and relapse rates remain unsatisfactory, underscoring the critical need for adjunctive and mechanistically novel treatments^5^.

While current pharmacotherapies primarily target the mu-opioid receptor, accumulating evidence suggests that glutamatergic signaling plays a central role in the neurobiological adaptations underlying opioid dependence and withdrawal^6–8^. N-methyl-D-aspartate receptors (NMDAR) have been targeted for chronic pain management, yet less emphasis has been placed on compounds that target this receptor for OUD^9,10^. NMDAR play a central role in synaptic plasticity, learning, and memory, all of which are dysregulated in addiction and withdrawal states. Moreover, chronic opioid exposure leads to dysregulation of NMDA receptor-mediated synaptic plasticity in key brain regions such as the nucleus accumbens and extended amygdala that significantly contributes to both the rewarding and aversive properties of opioids^8,11^.

Spiro-β-lactam-based NMDA positive allosteric modulators have shown promise in enhancing cognitive performance and inducing synaptic plasticity in preclinical models^12,13^. NYX-783 is a novel spirocyclic-β-lactam compound that mimics the dipyrrolidine-based β-turn motif found in rapastinel, a well-studied NMDA receptor modulator, and binds to a non-competitive site on the NMDAR distinct from the glutamate and glycine binding domains. Prior studies with related compounds have demonstrated improvements in learning and memory, as well as modulation of synaptic plasticity^14^. These properties suggest that NYX-783 and similar modulators may offer therapeutic advantages by supporting cognitive recovery and neuroadaptation in the context of OUD. Importantly, rapastinel (GLYX-13) and related modulators such as NYX-2925 have already been evaluated in clinical or translational contexts, including depression, PTSD, and neuropathic pain, establishing the conditions for repurposment of NMDA modulators toward OUD pharmacotherapy^15^. Yet, NYX-783 has not been assessed for its ability to remediate OUD-related conditions.

In this study, we investigated the effects of NYX-783 in a range of preclinical tests relevant to OUD to evaluate its potential as a candidate therapeutic. These included: (1) oxycodone-induced respiratory depression; (2) oral oxycodone self-administration and post-abstinence reinstatement to model relapse; (3) self-administration on a progressive ratio schedule to assess motivation for drug-seeking; (4) somatic withdrawal signs following precipitated withdrawal from oxycodone or morphine; and, (5) conditioned place aversion (CPA) as a proxy for negative affective states during withdrawal.

## Results

### NYX-783 does not exacerbate oxycodone-induced respiratory depression

First, we sought to establish safety for use of NYX-783 in the context of opioid exposure. Male and female mice had their respiratory rate monitored under increasing doses of oxycodone injected every twenty minutes. As expected, respiratory rate gradually decreased for both sexes over time as a result of the escalating oxycodone treatments^16^. No dose of NYX-783 (0, 0.1, 1.0, or 10.0 mg/kg ip) was observed to alter respiratory depression at any time point for either sex (Fig. 1A and B).

**Fig 1:**
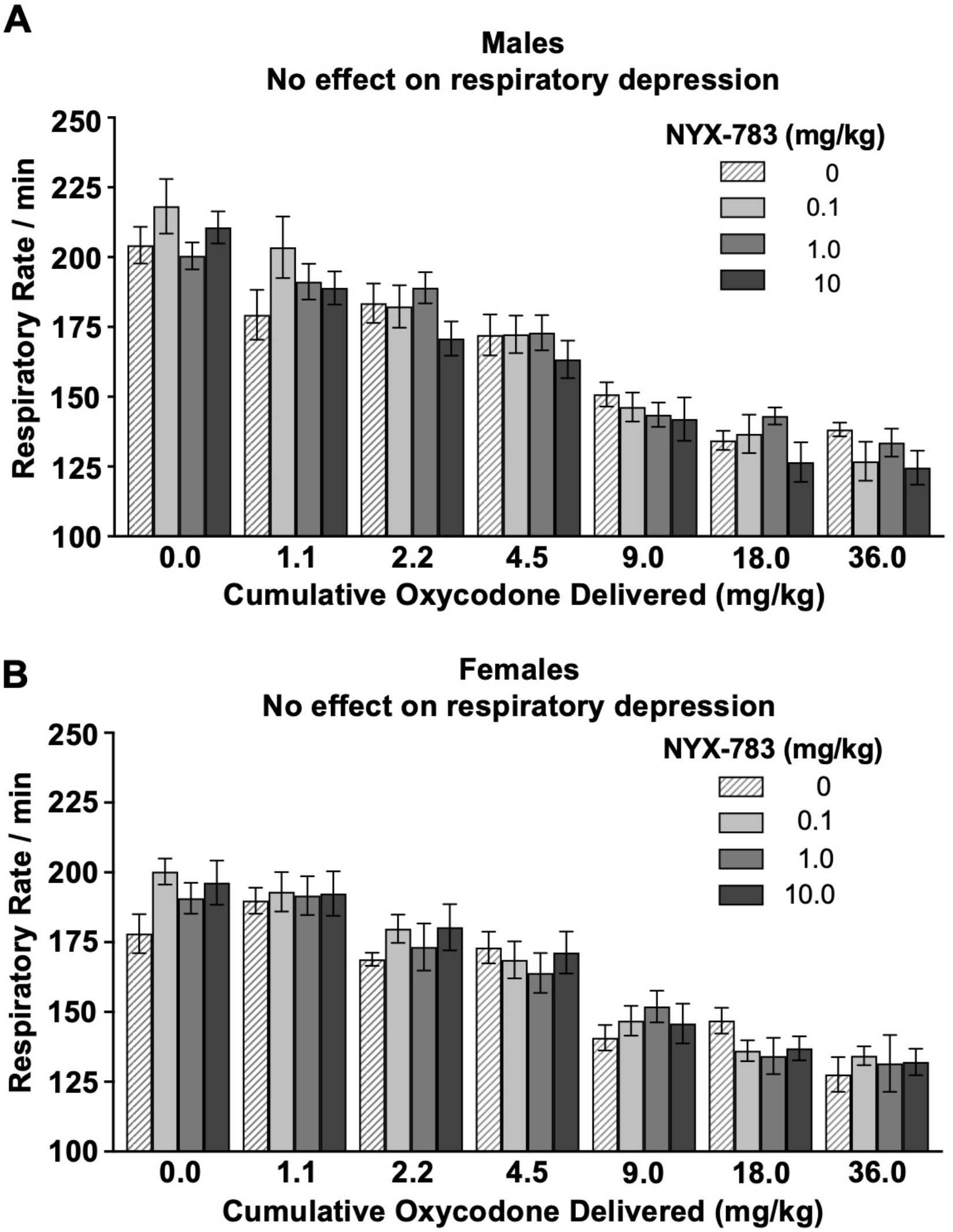
No effect of NYX-783 on oxycodone-induced respiratory depression in male or female mice. Respiratory rates per minute generally decreased as cumulative oxycodone increased. Mice were tested over a range of NYX-783 pretreatment doses. Separate cohorts were used for each dose of NYX-7683. **(A)** Male mice (n= 6-11) showed no effect of NYX-783 (at any dose) on opioid-induced respiratory depression. **(B)** Female mice (n = 6-8) showed no effect of NYX-783 (at any dose) on opioid-induced respiratory depression. All error bars are S.E.M.

### NYX-783 does not change oral oxycodone self-administration during maintenance, post-abstinence, or progressive ratio seeking

Male and female mice were trained to self-administer oral oxycodone in operant chambers with a previously established saccharin fade protocol^17^. All mice exhibited a typical profile of elevated self-administration of oxycodone when saccharin was included in the reinforcer solution, followed by a decrease during saccharin fading, and finally stability of intake during the maintenance phase after saccharin has been completely removed (Fig. 2A). After five days of maintenance self-administration, male mice were tested for effects of NYX-783 on oxycodone self-administration every other day in a Latin Square Design. No dose of NYX-783 was observed to alter self-administration of oxycodone (Fig. 2B). Separate cohorts of male and female mice were designated for post-abstinence testing. These mice underwent training for oral oxycodone in identical fashion, and then underwent abstinence for seven days after their maintenance phase. All mice were maintained *ad libitum* on standard chow during abstinence, and then food restricted the night prior to resuming self-administration and subsequent testing every other day in a Latin Square Design. No effect of treatment was observed for male or female mice (Figs. 2C, D). A separate cohort of female mice were trained and tested for effects of NYX-783 on self-administration on a PR schedule after their five day maintenance phase to evaluate motivation^18,19^. No effect of NYX-783 (1 mg/kg) was observed (Fig. 2E).

**Fig 2:**
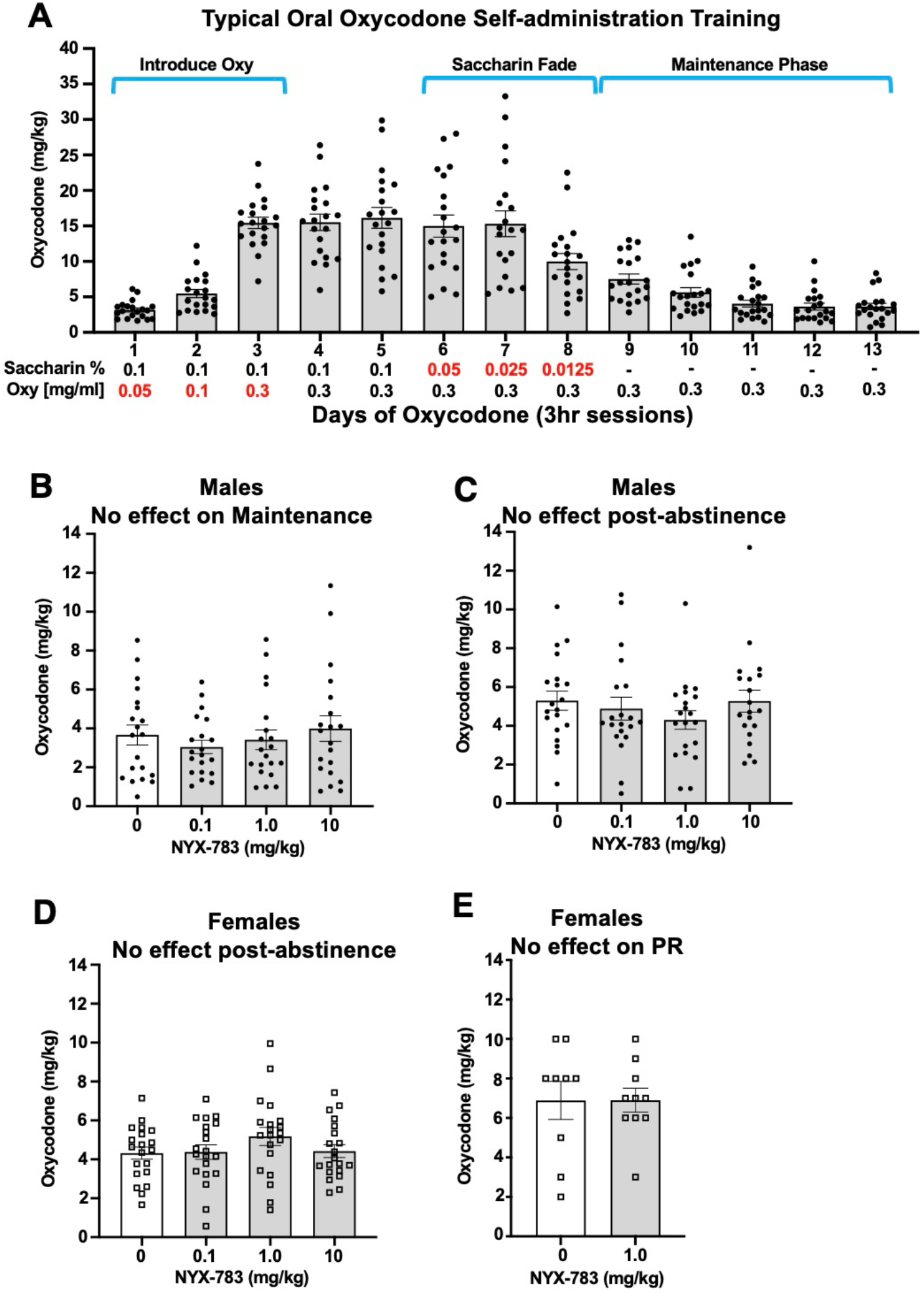
No effect of NYX-783 on different phases of self-administration for oxycodone. **(A)** A typical oral oxycodone training progression showing consumption across the various phases of training, ending with stability for consumption averaging ~ 3-5 mg/kg (males, n=20). **(B)** No effect of NYX-783 was observed in male mice (n=20) undergoing Latin Square design testing beginning on Day 14. **(C)** A separate cohort of male mice (n=20) were trained in identical fashion and then subjected to an abstinence period of seven days after their final maintenance Day 13 session. No effect of NYX-783 was observed upon self-administration post-abstinence. **(D)** Female mice (n=20) were trained and subjected to abstinence in an identical fashion. Latin Square testing showed no effects of NYX-783. **(E)** A separate cohort of females was trained and tested under a progressive ratio (PR) paradigm. NYX-7683 (1 mg/kg) had no effect on rewards earned (veh n=9, NYX-783 n=10). All error bars are S.E.M.

### NYX-783 has no effect on naloxone-induced withdrawal jumps

Male and female mice were used to assess NYX-783 on naloxone-precipitated withdrawal jumping behavior using morphine pellets or osmotic minipumps delivering oxycodone^20–22^. After five days of chronic exposure to the subcutaneous morphine pellet, mice were treated with NYX-783 (0, 0.1, 1.0, or 10 mg/kg) one hour prior to naloxone (1.0 mg/kg). Jumping behavior was immediately quantified over a 20-minute period after naloxone. No effect of NYX-783 was observed at any dose (Fig 3A, B). To verify the generalizability of these findings across opioids, osmotic minipumps were used to deliver chronic oxycodone to female mice. After three days of delivery, mice were treated with NYX-783 (0, 0.1, 1.0, or 10 mg/kg) one hour prior to naloxone (1.0 mg/kg). No effect of NYX-783 was observed at any dose (Fig 3C).

**Fig 3:**
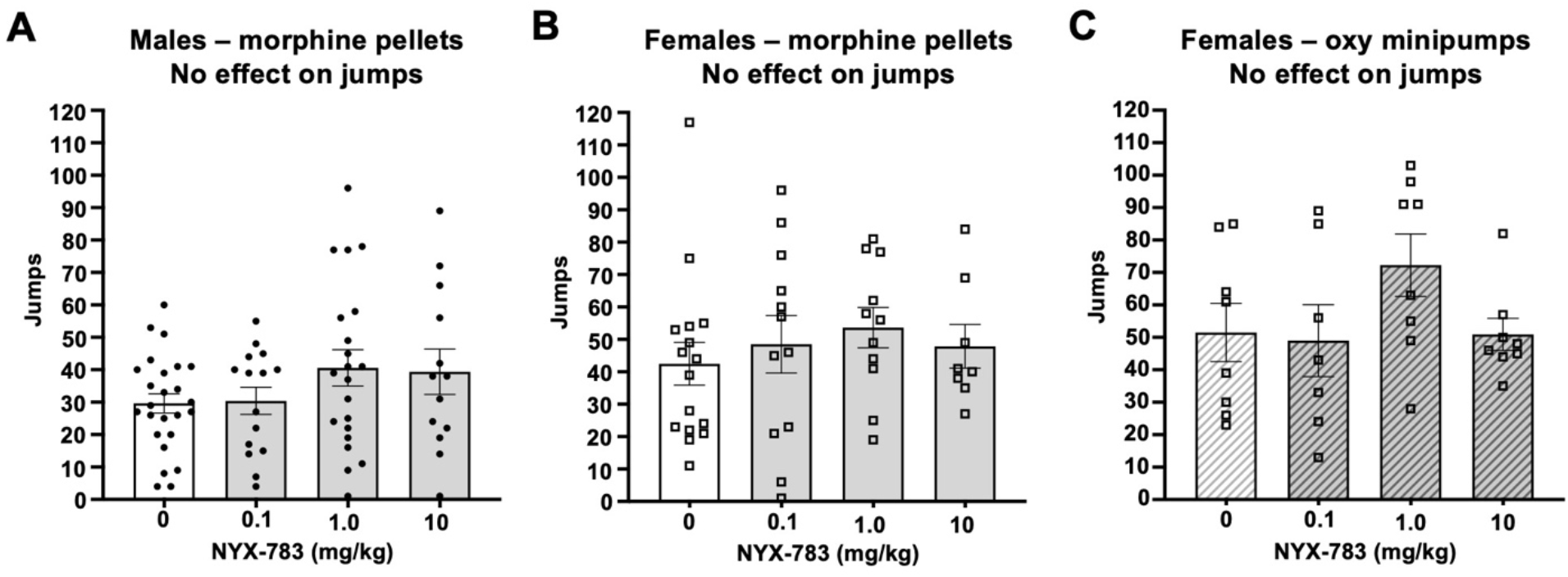
No effect of NYX-783 on somatic withdrawal jumps. **(A and B)** Male and female mice that were implanted with a morphine pellet were tested on Day 5 to assess somatic withdrawal jumps after naloxone-precipitated withdrawal. No effect of pretreatment with NYX-783 one hour prior to the naloxone was observed for any dose (n= 25, 15, 21, and 13 for 0, 0.1, 1.0, and 10 mg/kg doses respectively for males; and n= 16, 12, 11, and 8 for 0, 0.1, 1.0, and 10 mg/kg respectively for females). **(C)** No effect of NYX-783 was seen in female mice chronically administered oxycodone 130 mg/kg/day via osmotic minipump prior to naloxone (n= 8, 7, 8, 8 for 0, 0.1, 1.0, and 10 mg/kg doses NYX-783 respectively). All error bars are S.E.M.

### NYX-783 reduces aversion during withdrawal

Male and female mice were used to assess effects of NYX-783 on the aversive component of opioid withdrawal as measured by conditioned place aversion (CPA). All mice under went alternating days of aversion and neutral pairings over six days each. oxycodone-NYX-783-naloxone (aversion pairing) and saline-saline-saline (neutral pairing) in the respective pairing chambers. Aversion pairing days involved oxycodone followed by Veh or NYX-783 (10 g/kg) one hour later, followed by naloxone (0.1 mg/kg) one hour later and immediately prior to confinement in the aversion designated chamber. This dose of naloxone was not sufficient to reproduce the jumping behavior described above, but was sufficient to result in CPA, as confirmed by one sample t-tests. Separate cohorts of mice were used for Veh or NYX-783 treatments. On test day, mice were given free access to all chambers. NYX-783 treatment during aversion pairing resulted in a near-significant attenuation of withdrawal-induced CPA in males, and a significant reduction in females (Fig 4A, B).

**Fig 4:**
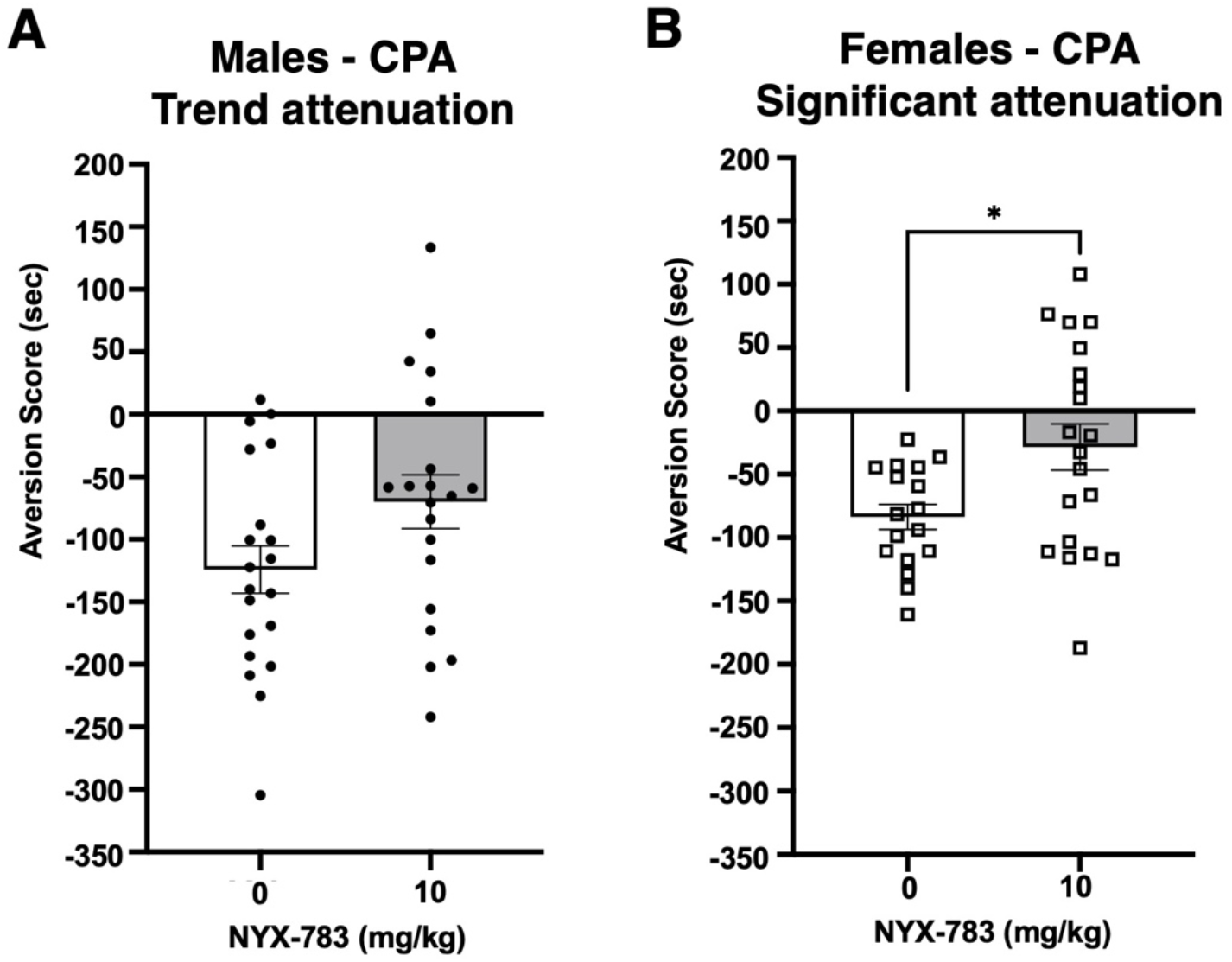
NYX-783 reduces opioid withdrawal-conditioned aversion. After three rounds of NYX-783 (10 mg/kg) pretreatment on aversion pairing days interspersed with neutral pairing days, mice were tested for the development of conditioned place aversion. **(A)** Male mice treated with NYX-783 showed a trending improvement in their aversion score compared to Veh mice (n=20, 20; Welch’s t-test, df=37.43, t=1.897, *p*=0.0656).To assess the effectiveness in generating conditioned aversion, a one sample t-test revealed a significant CPA score for the Veh treatment group (t=6.548, df=19 *p*<0.0001), as well as a significant CPA score for the NYX-783 group (t=3.254, df=19, *p*=0.0042) vs a theoretical mean of 0.00. **(B)** Female mice treated with NYX-783 showed a significant improvement in aversion score compared to Veh mice (n=17, 20; Welch’s t-test, df=28.80, t=2.668, \**p*=0.0124). To assess the effectiveness in generating conditioned aversion, a one sample t-test revealed a significant CPA score for the Veh group (t=8.500, df=16, *p*<0.0001), and no significant CPA score for the NYX-783 group vs a theoretical mean of 0.00. A 2-way ANOVA including sex and treatment revealed a significant main effect of sex (F_(1, 73)_ = 5.055, \**p*=0.0276) and a significant main effect of treatment (F_(1, 73)_ = 9.045, \*\**p*=0.0036), however since the male and female cohorts were collected separately, the interpretation of sex-specific effects is inconclusive at this time. All error bars are S.E.M.

## Discussion

The purpose of this study was to evaluate the potential use of the NMDAR positive allosteric modulator NYX-783 for treatment of OUD, using behavioral and physiological tests in mice. Our principal finding is that NYX-783 attenuated CPA to precipitated opioid withdrawal. This behavioral effect was robust in female mice, and in male mice a similar reduced pattern of CPA, with trending significance, was observed. Because CPA may be a validated preclinical proxy for dysphoric states that emerge during withdrawal, these data position NYX-783 as a candidate agent for clinical assessment for the alleviation of states of affective malaise that precipitates relapse in OUD^23–27^.

The ability of NYX-783 to attenuate CPA aligns with evidence that both NMDA receptor antagonists and positive modulators can disrupt the acquisition of opioid withdrawal aversion, likely by modulating glutamatergic signaling and synaptic plasticity in the nucleus accumbens and amygdala circuits^8,11^. Furthermore, recent preclinical work has highlighted sex differences in the neurobiological response to opioid withdrawal, with females showing distinct patterns of synaptic plasticity and affective processing in reward-related brain regions, which may contribute to the clearer effects of NYX-783 observed here in female animals^28,29^. While we are unable to directly compare males and females in this study, higher sensitivity in females is consistent with a broader literature on sex differences in addiction models and withdrawal responses^30,31^, and suggest that positive allosteric modulation of NMDA receptors may selectively target the negative affective and motivational symptoms of opioid withdrawal.

Notably, NYX-783’s reduction of aversion was not complemented by alterations in somatic withdrawal, assessed here by naloxone-precipitated jumping. No significant changes in jumping behavior were detected in either sex for either method, suggesting that NYX-783 does not alleviate or worsen physical withdrawal symptoms. The ability of NYX-783 to attenuate negative affective symptoms of opioid withdrawal without altering somatic signs may be therapeutically advantageous; Clinical literature highlights that affective symptoms like anxiety, dysphoria, and craving can persist much longer than somatic withdrawal signs and are primary drivers of relapse risk^6,25,26^. A compound that mitigates negative affect without dampening somatic feedback avoids the risk of obscuring physiological withdrawal severity, thereby complementing existing pharmacotherapies. In this way, NYX-783 appears to act with specificity on the affective dimensions of OUD, rather than exerting broad suppressive or nonspecific effects.

Given that opioid-induced respiratory depression is the primary cause of opioid-related mortality, we also assessed whether NYX-783 exacerbated this outcome using a paradigm of escalating oxycodone doses with concurrent monitoring of respiratory rate^16^. Pretreatment with NYX-783 had no impact on oxycodone-induced respiratory depression in either male or female mice across multiple doses. This result satisfies a critical safety milestone, particularly given that many existing OUD medications, including full and partial opioid agonists, carry a major risk of inducing respiratory depression. Prior work has extensively characterized the mechanisms of opioid-induced respiratory depression and highlighted potential non-opioid strategies to mitigate it^32^, making it essential to evaluate new therapeutic candidates in this context. The absence of any potentiation of this adverse effect by NYX-783 supports its development as a non-opioid adjunctive therapy, and complements reported positive safety findings of rapastinel and related NMDA modulators in clinical trials.

Operant self-administration assays—encompassing maintenance intake, post-abstinence relapse, and progressive-ratio (PR) motivation tests—showed no effect of NYX-783 on oxycodone consumption or drug-seeking behavior. This is consistent with prior evidence that NMDA receptor modulation does not universally suppress opioid reinforcement mechanisms, but may instead act on specific neurobiological substrates of withdrawal and negative affect^11^. The apparent specificity of NYX-783 for aversive withdrawal states, rather than general reward suppression, reduces concerns about anhedonia or interference with natural reward processing– a limitation of some other pharmacological interventions for OUD.

Taken together, our findings suggest that positive allosteric modulation of NMDA receptors can selectively target the negative-affective symptoms of opioid withdrawal without altering somatic withdrawal, respirat, or drug-taking behavior. These results represent a conceptual shift from mu-opioid receptor–centric approaches toward interventions that normalize glutamatergic dysregulation and synaptic plasticity. Targeting the affective state of addiction and dependence is increasingly recognized as a necessary complement to existing agonist or antagonist therapies, as these states are key contributors to seeking and relapse^24–26^. From a translational perspective, the lack of effect of NYX-783 on oxycodone self-administration also reduces concern about abuse liability or interference with natural rewards, while the absence of respiratory potentiation supports safety. Future work should also test NYX-783 in combination with existing medications such as buprenorphine or extended-release naltrexone, to determine whether adjunctive strategies further reduce aspects of relapse liability. Collectively, these results position NYX-783 and related spirocyclic modulators as promising candidates for next-generation OUD therapeutics aimed at alleviating the affective states that drive relapse.

### NYX-783 Methods

#### Animals

Animal experiments were carried out in accordance with Yale University School of Medicine Institutional Animal Care and Use Committee (IACUC) regulations. All protocols were approved by the Yale IACUC, and all surgeries were performed in compliance with approved pre- and post-operative care. All methods are presented in accordance with ARRIVE guidelines with the exception that the experimenters were not blinded to the treatments. Animals removed from the experiment included: 1) three female mice from the CPA experiment due to accidental pairing with the incorrect chamber, 2) four males that had 0 naloxone-induced jumps (*a priori* criteria), and 3) one female undergoing oral oxycodone self-administration for the progressive ratio experiment that failed to consume oxycodone rewards earned. Male and female C57BL/6J mice, ages 8-18 weeks were used for these studies (Jackson Laboratories, Bar Harbor). Mice were group housed up to five per cage and maintained *ad libitum* with standard chow (Teklad Global #2018) on a 12-hour light/dark cycle (lights on at 7 a.m.). For self-administration studies, mice were food restricted and maintained at 85-90% of their initial bodyweight.

#### Drugs and reagents

NYX-783 was provided by Aptinyx Inc. and dissolved in sterile saline, aliquoted, and stored at −20C. Saline was used as the vehicle. Oxycodone HCl was provided by the NIDA Drug Supply Program. Saccharin (Acros Organics #149005000) was diluted to several concentrations in tap water for self-administration experiments. Morphine pellets (25 mg morphine sulfate) were provided by the NIDA Drug Supply Program. Osmotic minipumps (Alzet 1003D) were custom loaded to deliver 130 mg/kg/day oxycodone. Naloxone HCl was purchased from Spectrum Chemical (#N1231) and dissolved in sterile saline. Lidocaine (Covetrus, North America VINB-0024-6800) was diluted with sterile saline to deliver 4 mg/kg subcutaneously at the incision site for subcutaneous morphine pellet implantation and osmotic minipumps. Isoflurane (Covetrus) was used to anesthetize mice for subcutaneous surgeries and hair removal.

#### Statistics

ANOVAs were used to assess statistical significance for Latin Square Design experiments. A mixed model ANOVA was used to analyze respiratory depression data. Welch’s t-test (two-tailed) was used to assess significance for the withdrawal jumps and CPA studies. One sample t-tests were used to assess the effectiveness of the CPA paradigm. All analyses were conducted using Prism (GraphPad Software), and the statistical significance threshold was set at p < 0.05.

#### Respiratory depression

Young adult male and female C57BL/6J mice were used. Respiratory rates were assessed using the MouseOx Plus System (Starr Life Sciences). A 12.5 mm 300 rpm capsule electrical slip ring (Comidox) was spliced into the MouseOx Plus collar wires to reduce torsional strain caused by ambulatory activity of the mouse. The day prior to testing, mice underwent neck hair removal with Nair while briefly anesthetized with isoflurane. On the test day, mice were habituated to the MouseOx Plus collars (Starr Life Sciences) for one hour prior to injections. Mice were then pretreated with NYX-783 (0.1, 1.0, 10.0 mg/kg or saline ip) one hr prior to escalating doses of oxycodone. Oxycodone (0, 1.1, 1.1, 2.2, 4.5, 9.0, and 18.0 mg/kg ip) was delivered every 20 minutes^16^. Respiration was monitored at 5 Hz over a 15-minute period immediately after each oxycodone injection. Opioid-induced locomotor activity occasionally disrupted data acquisition, resulting in variability of the sample size (n) across treatment conditions. Separate cohorts of mice were used for each dose of NYX-783.

#### Operant self-administration and progressive ratio

Male and female mice were food restricted and maintained at 85-90% of their starting bodyweight throughout training and self-administration^17,33^. Med Associates mouse operant boxes with syringe pumps were used for self-administration sessions. The boxes consisted of three nose poke ports on one wall, and a magazine for liquid reward delivery on the opposite wall. Infrared beams were used to monitor nose entries in each nose poke port. Cue lights were used to indicate the nose poke port activation, as well as the reward delivery in the magazine. 5 ml syringes were loaded with the rewards, and the pump activation time was set to deliver a reward size of 20 µl.

Mice first underwent at least two 30-minute magazine training sessions on different days for a 0.1% saccharin solution delivered to the magazine without any operant requirement. Next, mice were trained to actively nose poke for each reward on a fixed ratio schedule of one nose poke per reward (FR1). The cue light in the active nose poke port was turned on when the port was initially activated by the program, and upon successful nose entry was turned off. The reward was delivered immediately, and the cue light in the magazine was turned on to indicate the presence of a reward. Upon entry to the magazine, that cue light was turned off and after a 5 sec delay, the nose poke port and cue light became active again. After the mice learned to operantly self-administer the 0.1% saccharin solution on the FR1 schedule, the nose poke response requirement was increased to three nose pokes per reward (FR3 schedule). After several days, oxycodone was gradually introduced to the saccharin solution, starting at 0.05 mg/ml and increased to 0.3 mg/ml, after which saccharin was faded out over four days. Five additional ‘maintenance days’ of self-administration after the saccharin fade were performed to evaluate stability in operant behavior and consumption of unsweetened oxycodone. Magazines were inspected after each session to verify consumption, and mice that consistently left oxycodone behind were excluded.

NYX-783 was tested using a within-subject counterbalanced Latin Square Design in which each mouse received all doses (0.1, 1.0, 10 mg/kg, or saline ip) one hour before its self-administration session for 0.3 mg/ml unsweetened oxycodone, with test days separated by two additional maintenance days. In separate animals, instead of initiating NYX-783 challenge directly following the initial maintenance self-administration period, a ‘post-abstinence’ self-administration procedure was followed. Here, mice underwent a seven-day abstinence period (ad libitum chow, no operant sessions), after which they were again food-restricted the night prior to resuming self-administration, and tested in the Latin Square protocol.

A separate cohort of female mice were trained to self-administer oxycodone on the same procedure as above, followed by testing on a progressive ratio (PR) schedule with an increasing nose poke response requirement per reward (nose poke requirements 1, 2, 4, 6, 9, 12, 15, 20, and 25 for the first ten rewards). Prior to testing, mice were assigned to Vehicle (saline) or NYX-783 (1.0 mg/kg ip) groups, counter-balanced by their last three days of self-administration performance. Due to the relatively short half-life of NYX-783 (less than five hours), no break point was set *a priori*, and thus mice being treated one hour prior to their sessions were allowed to nose poke for the next reward throughout the three-hour test session.

#### Somatic withdrawals

Male and female mice were used to measure naloxone-induced withdrawal jumps. Med Associates locomotor boxes with infrared (IR) beams were used to assess jumps. The beam boxes were elevated (approximately 6 inches) such that the IR beams could only be broken when the mouse jumped with all four paws leaving the cage bottom, and specifically not broken due to rearing or partial jumps. A 0.5-second timeout was added to the custom software program (Med-PC V) after the event of a single beam break to avoid double-counting from a single jump. Clear plastic mouse buckets with padded high-profile lids to accommodate jumping behavior were used. Data was collected with Med PC V software.

Mice were anesthetized with isoflurane (2-3%), injected with lidocaine at the implantation area, shaved, and disinfected with betadine prior to incision. After a small incision, a single 25 mg morphine pellet was inserted subcutaneously, and the incision was closed with wound clips and coated with triple antibiotic ointment before returning them to their housing room. On the fifth day after implantation, mice were pretreated with NYX-783 (0.1, 1.0, 10 mg/kg or saline ip) one hour prior to naloxone (1 mg/kg ip). Jumping behavior was immediately quantified over the next 20 minutes. Osmotic minipumps were custom loaded to deliver 130 mg/kg/day. On Day 3, mice were pretreated with NYX-783 (0.1, 1.0, 10 mg/kg or saline ip) one hour prior to naloxone (1 mg/kg ip), and jumping behavior was immediately quantified over the next 20 minutes.

#### Conditioned place aversion

CPA studies were conducted in Med Associates 3-chambered boxes, consisting of two distinct conditioning chambers separated by a smaller neutral gray chamber with guillotine doors. Mouse movement and position throughout the box were tracked with infrared beams. On Day 1 (habituation), mice were placed in the middle gray chamber and given free access to both conditioning chambers for fifteen minutes. Time spent in the two conditioning chambers was monitored, and then used to assign aversion pairing to the chamber that the mouse spent the most time in. On days 2, 4 and 6 mice went through aversion pairing where they received an acute dose of oxycodone (5 mg/kg ip, 2 hours prior to pairing) and of NYX-783 (10 mg/kg or saline ip, 1 hour prior to pairing) followed by naloxone (0.1 mg/kg ip, immediately prior to pairing). Mice were then confined to the assigned conditioning chamber (pairing) for 30 minutes. On days 3, 5, and 7, neutral pairings were conducted with three saline (ip) injections at 2 hours,1 hour, and finally immediately prior to confinement in the opposite chamber for thirty minutes. On day 8 (test day), mice were placed in the middle chamber and allowed free access to all pairing chambers over fifteen minutes. Aversion scores were calculated as the time in the aversion chamber on test day minus the time spent in that chamber on habituation day (A = Time_test_ – Time_habituation_).

## Acknowledgements

Work was supported by the NIH (UG3DA050322). This work was also funded in part by the State of Connecticut, Department of Mental Health and Addiction Services, but this publication does not express the views of the Department of Mental Health and Addiction Services or the State of Connecticut. The views and opinions expressed are those of the authors. We would also like to acknowledge the role of Dr. Ronald Duman in encouraging and enabling the initiation of this work.

